# Blunted response of caudal locus coeruleus to arousing stimuli in Parkinson’s Disease

**DOI:** 10.1101/2025.08.13.670094

**Authors:** Anders E. Lund, Christopher F. Madelung, Kristoffer H. Madsen, Birgitte L. C. Thomsen, Annemette Løkkegaard, David Meder, Hartwig R. Siebner

## Abstract

Parkinson’s disease (PD) causes progressive degeneration of noradrenergic neurons in the locus coeruleus (LC), contributing to non-motor symptoms. Using neuromelanin-sensitive ultra-high field (7T) MRI, we previously identified a reduction in neuromelanin signal in the caudal LC, indicating a rostro-caudal gradient of noradrenergic cell loss. Caudal LC degeneration was associated with greater severity of non-motor symptoms such as orthostatic hypotension and apathy.

In the current study, we expanded the PD cohort to further validate the structure–symptom relationships within the LC and investigate how degeneration along the rostro-caudal LC axis affects arousal-related functional responsivity. To this end, 71 people with PD in the ON-medication state and 40 age- and sex-matched healthy controls underwent clinical assessments and 7T magnetization transfer-weighted (MTw) MRI to quantify structural changes along the rostro-caudal LC axis. A subgroup of 30 people with PD and 27 controls underwent 7T fMRI to assess LC responsivity to arousing auditory and visual stimuli in two fMRI sessions on separate days. Healthy controls were scanned twice without medication, while people with PD were studied on and off dopaminergic medication in counterbalanced order.

In the PD group, MTw MRI confirmed a significant reduction of the regional neuromelanin signal in caudal LC relative to the control group (*P* = 0.0099). This structural disintegration correlated with orthostatic hypotension (*P* = 0.0087) and cognitive impairment (*P* = 0.036), corroborating its clinical relevance. Functional MRI revealed reduced activation of the caudal LC to arousing visual and auditory stimuli in people with PD relative to controls (*P* = 0.012). This difference reached statistical significance only in the ON-medication state, with a similar but non-significant trend in the OFF-medication state (*P* = 0.10). In an exploratory analysis of a smaller sub-sample, structural and functional caudal LC signals were significantly correlated in both people with PD and healthy controls (*P* = 0.0069).

Together, the findings provide evidence for a rostro-caudal gradient of LC pathology in PD at both structural and functional levels. While structural MRI provides fine-grained insights into spatial gradients of disease-related pathology, functional MRI captures impaired functional responsivity of caudal LC. The presence of arousal-induced hypoactivation of caudal LC in the ON-medication state indicates that LC dysfunction extends beyond dopamine deficits in PD, highlighting complex interactions between dopaminergic and noradrenergic systems.

## Introduction

Parkinson’s disease (PD) is a multi-system neurodegenerative disease characterized by progressive loss of vulnerable neurons. The neurodegenerative process also includes the locus coeruleus (LC).^1^ 20-90% of its neurons degenerate during the course of PD, with neuronal loss often preceding and exceeding dopaminergic degeneration.^2,3^

The LC is the brain’s primary source of noradrenaline, containing 30.000-50.000 noradrenergic neurons. Noradrenaline release plays a central role in regulating arousal,^4^ characterized physiologically by sympathetic nervous system activation.^5^ The noradrenergic neurons in LC become pigmented by neuromelanin in mid and later life and its dark pigmentation from neuromelanin makes it visually distinct in postmortem brain tissue.^6–8^ The LC sends noradrenergic projections throughout the brain, modulating functions such as arousal, sleep, emotion, executive function, and autonomic regulation.^4,9^ Consequently, LC degeneration contributes to non-motor symptoms, such as apathy, cognitive impairment, sleep disturbances, and orthostatic hypotension,^10,11^ which often burden patients more than motor symptoms.^12,13^ LC degeneration also impairs arousal responses, leading to secondary effects on cognition, sleep, and bodily functions.

The LC is a small paired brainstem nucleus with a thin, spindle-like shape (∼15 × 2 × 2 mm) along the floor of the fourth ventricle in the rostral pons.^14^ Its small diameter (2 x 2 mm) makes in vivo MRI of its structure and function a challenge. Neuromelanin sensitive MRI sequences have enabled visualisation of noradrenergic neuron density as a marker of structural LC integrity.^1,8,15^ Both, neuromelanin sensitive MRI and histological studies, indicate that LC degeneration in PD is predominantly in the caudal region.^16,17^ Since task-based functional studies of LC responses are lacking,^18^ it remains to be examined how LC degeneration alters its functional responses in PD. We and others have previously shown associations of structural disintegration in the LC with clinical symptoms.^16,19,20^ However, whether the individual degree of impairment in functional responses of the LC is associated with non-motor symptoms is yet to be explored.

This study adopted a multimodal 7T MRI approach to identify structural and functional LC alterations along its rostro-caudal axis in PD and link these alterations to non-motor symptoms. The study had two primary aims: First, we employed 7T fMRI to assess sub-regional LC responses to arousal stimuli in an auditory and a visual event-related fMRI paradigm. In line with our pre-registered hypothesis, we tested whether people with PD would show attenuated LC activation in response to these stimuli. Second, we extended our previously reported structural MRI dataset to further characterize LC degeneration in PD.^16^ We hypothesized that neuromelanin sensitive 7T MRI would reveal selective caudal LC degeneration. We expected caudal LC degeneration to correlate with clinical markers of noradrenergic dysfunction, specifically systolic drops in orthostatic blood pressure and apathy,^16,17^ as well as cognitive impairment.^19–22^ We further hypothesized that reduced functional LC activation would correlate with structural disintegration.

## Materials and methods

### Participants

Structural MRI and non-motor symptoms were assessed in 114 participants, comprising 74 people with idiopathic PD and 40 healthy, age- and sex-matched healthy controls (HC) (Table 1). Amongst these are 42 people with PD and 24 HC that we have described earlier in terms of structural change in the LC.^16^ People with PD were recruited via the Movement Disorders outpatient clinic at the Department of Neurology, Copenhagen University Hospital Bispebjerg (Copenhagen, Denmark), private practice neurology clinics in the Copenhagen Region, and online advertisement. Inclusion criteria for the PD group were (I) a clinical diagnosis of PD assessed by a neurologist and (II) meeting the MDS Clinical Diagnostic Criteria for PD.^23^ Exclusion criteria were pregnancy or breastfeeding, history of other neurologic or psychiatric disease, implanted electronic devices, and claustrophobia. Three people with PD were excluded due to poor scan quality. HC were recruited through online advertisement or as relatives to patients with PD. They were required to be ≥18 years old, meeting none of the exclusion criteria mentioned above, and having no history of significant neurologic or psychiatric disease.

**Table 1.** Demographic and clinical characteristics of the population.

|  | Structural MRI (n = 111) |  |  | Arousal study subgroup (n = 57) |  |  |
| --- | --- | --- | --- | --- | --- | --- |
|  | All people with Parkinson's disease (n = 71) | All healthy controls (n = 40) | P | People with Parkinson's disease (n = 30) | Healthy controls (n = 27) | P |
| Age (years) | 67.0 (38.0-88.0) | 63.0 (40.0-80.0) | 0.089 | 60.5 (38.0-79.0) | 59.0 (40.0-80.0) | 0.92 |
| Sex (male/female) | 44/27 | 25/15 | 1 | 19/11 | 17/10 | 1 |
| Disease duration (years) | 3.93 (0.250-19.2) | - | - | 3.34 (0.260-11.2) | - | - |
| Time since symptom onset (years) | 6.06 (0.980-21.3) | - | - | 4.61 (1.07-13.9) | - | - |
| Most affected side (left/right) | 33/38 | - | - | 14/16 | - | - |
| Systolic orthostatic blood pressure change | -16.0 (-72.0,17.0) | - | - | -8.50 (-70.0,17.0) | - | - |
| UPDRS-III (mean) | 26.3 (10.5) | - | - | 22.4 (7.42) | - | - |
| NMSS | 32.0 (3.00-138) | - | - | 27.0 (3.00-113) | - | - |
| MoCA | 27.0 (17.0-30.0) | 28.0 (23.0-30.0) | 0.075 | 28.0 (23.0-30.0) | 28.0 (25.0-30.0) | 0.53 |
| BDI-II | 6.00 (0.00-20.0) | 3.00 (0.00-20.0) | 0.00013 | 5.00 (0.00-14.0) | 3.00 (0.00-20.0) | 0.0099 |
| LARS (mean) | -24.8 (6.04) | -26.8 (4.96) | 0.061 | -26.5 (4.99) | -27.8 (4.21) | 0.31 |

Data are presented as median unless otherwise indicated. Comparisons of patient and control groups were assessed using independent t-tests for normally distributed data and Wilcoxon rank-sum test when normal distribution could not be assumed. For sex distribution and handedness between-group comparisons were assessed with Pearson's chi-squared test with Yates' continuity correction and Fisher's exact test, respectively. Abbreviations: UPDRS-III = Unified Parkinson's Disease Rating Scale Part III; NMSS = Non-Motor Symptom Scale; MoCA = Montreal Cognitive Assessment; BDI-II = Beck's Depression Inventory II; LARS = Lille Apathy Rating Scale.

For assessing the functional arousal changes in the noradrenergic system, a subgroup of 30 people with PD and 29 HC were tested on two additional test days (Fig. 1A). The inclusion criteria for this group were the ability to lie still during longer scan sessions, no known hearing loss, audiometric thresholds within 40 dB of normal hearing level and symmetric audiograms, defined as less than 40 dB difference between ears at two or more neighbouring frequencies. Further, people with PD had to pause their anti-parkinsonian medication for >12 hours for levodopa and >48 hours for other PD medication (e.g. dopamine agonists, amantadine, MAO-B inhibitors and COMT inhibitors) for one of the test days. Two HC were excluded due to poor scan quality.

**Figure 1.**
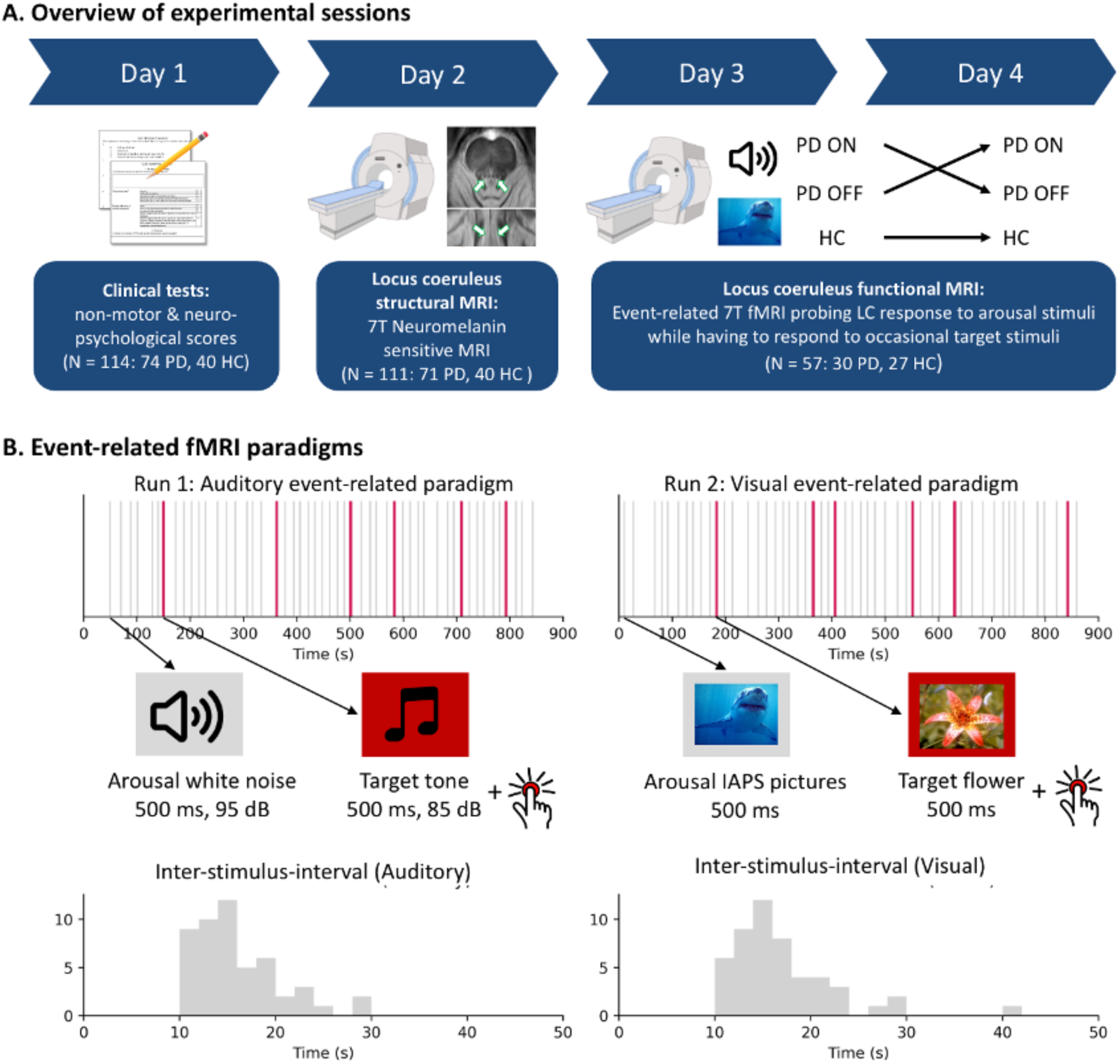
Study Design. **(A)** Overview of experimental sessions. The multimodal approach aimed to map noradrenergic changes in PD measuring non-motor symptoms, LC degeneration, and LC dysfunction. These were measured on four different sessions. The first session consisted of clinical tests, including neuropsychological tests, and assessment of non-motor symptoms. The second session consisted of structural ultra-high field (7T) neuromelanin sensitive MRI. The third and fourth sessions assessed noradrenergic function of the LC with two arousal paradigms during 7T fMRI. PD patients performed one session of arousal paradigms on medication and one session off medication, in randomized order. Healthy controls performed both sessions in the same state. **(B)** Event related fMRI paradigms. The auditory event-related paradigm (to the left) consisted of 45 white noise bursts (grey) played at 95 dB for 500 ms at pseudorandom intervals. Six times during the paradigm participants heard a target tone (red) for which they were instructed to press a button. The visual paradigm (to the right) had a similar design, showing 45 different arousing IAPS pictures (grey) for 500 ms. Six times participants were shown a flower (red) for which they were instructed to press. Each paradigm lasted 14 minutes and between stimuli participants were asked to gaze at a fixation cross on the screen. The inter-stimulus-interval was at least 10 s. Abbreviations: PD = Parkinson’s Disease; HC = Healthy controls; LC = Locus Coeruleus; 7T = 7 Tesla; fMRI = functional magnetic resonance imaging; dB = decibel; IAPS = International Affective Picture System.

The study was approved by the Regional Committee on Health Research Ethics of the Capital Region of Denmark (Record-id: H-18021857). All participants provided written informed consent to participate in the study in accordance with the Declaration of Helsinki. The study was pre-registered at ClinicalTrials.gov [NCT03866044].

### Study procedure

The first experimental day consisted of different examinations, including a neurological examination, Montreal Cognitive Assessment (MoCA)^24^, Barratt Impulsiveness Scale (BIS-11)^25^, Lille Apathy Rating Scale (LARS)^26^, Beck’s Depression Inventory (BDI-II)^27^, and Edinburgh Handedness Inventory^28^ (Fig. 1A). Motor and non-motor symptoms in people with PD were assessed in the ON-state using the Unified Parkinson’s Disease Rating Scale (UPDRS)^29^ and the Non-Motor Symptom Scale^30^. To assess orthostatic hypotension, patients’ blood pressure was measured after five minutes lying in a supine position. After standing up, several measurements were taken over a 3-minute period and the maximum change in systolic blood pressure was recorded.

On experimental day 2, participants underwent neuromelanin-sensitive structural 7T MRI. To assess eligibility for the functional sessions, an audiogram was performed (see Supplementary Methods).

A subgroup of participants performed a third and fourth session on two separate experimental days. On each day, participants arousal response was assessed once with fMRI and once psychophysiologically outside the scanner (to be reported separately). People with PD performed one session in the ON- and one in the OFF-state (counterbalanced). HC also underwent two sessions to balance session effects, but without medication. These sessions were performed at a similar time of day to balance circadian changes in noradrenaline levels.

### Event-related arousal paradigms

We designed one auditory and one visual arousal eliciting event-related paradigm (Fig. 1B). These paradigms was inspired by previous work^31^ and comparable stimuli have been shown to elicit arousal responses and LC activity.^31–34^

The auditory paradigm consisted of 45 bursts of white noise for a duration of 500 ms (see Supplementary Methods).^31^ Auditory stimuli were presented with pseudorandomized inter-stimulus-intervals (ISIs) sampled from a shifted exponential distribution, thus creating a constant hazard rate, i.e. a constant conditional probability of stimulus occurrence over time given that it has not occurred yet. Additionally, during the experiment, participants heard six target tones (500 ms, Supplementary Methods), to which they had to respond with a button press to keep them activated and listening. The sounds were delivered through Sensimetrics S15 in-ear-headphones which were isolated with self-hardening paste and foam cushions to minimize scanner noise. Participants were instructed to fixate on a cross hair to minimize eye movement related activity in the brainstem.

The visual paradigm (presented with PsychoPy2) had a similar structure and consisted of 45 stereotypically dangerous animals from the International Affective Picture System (IAPS) with high arousal and negative valence (pleasure-displeasure) ratings.^35^ The images were shown for 500 ms and in between participants were instructed to fixate on a cross hair. Similar to the noise paradigm, participants had to respond to six interspersed target images of a flower. If required, participants were offered MR-compatible glasses.

### Magnetic resonance imaging data

All MRI data were acquired on a Philips Achieva 7T scanner (Philips Best, Netherlands) with a 32-channel head coil (Nova Medical, Wilmington, MA, USA).

Structural MRI data were collected following an in-house protocol.^16^ Patients were scanned in the ON-state. First, to guide co-registration and normalization, we acquired whole-brain T1-weighted (T1w) Magnetization Prepared RApid Gradient Echo (MPRAGE) images (voxel size: 1 mm isotropic). Second, as a measure of LC integrity, we acquired magnetization transfer-weighted (MTw) images using a 3D high-resolution, ultra-fast gradient echo sequence with saturation pre-pulses aligned to the AC-PC line with a FOV covering the midbrain and rostral pons (voxel size 0.4 x 0.4 x 1.0 mm). For details about the sequences see Supplementary Methods.

During the fMRI sessions, we first acquired a structural 1.0 mm isotropic T1w whole-brain image for planning and co-registration. Second, B0-shimming was used to decrease the field inhomogeneity. For fMRI, we used EPI with a reduced transverse FOV covering the LC and midbrain aligned to the AC-PC line (voxel size: 1.0 x 1.0 x 2.2 mm, echo time = 23 ms, repetition time 1.2 s, number of slices: 20, slab size: 33 mm). A reverse phase-encoding image was obtained for distortion correction. Pulse oximetry and respiration were acquired for denoising.

### Structural image processing

Image processing, co-registration, and signal extraction of the MTw images followed an in-house pipeline from Madelung et al. using Advanced Normalization Tools (ANTs) software,^16^ which closely resembles the pipeline from recent publications.^19,36,37^

MT-weighted and T1-weighted images were bias field corrected.^38^ Within-subject co-registration and normalization to template space was performed by rigid-body, affine, and non-linear registration. First, to evaluate registration accuracy, high resolution images were transformed to individual subjects’ T1w space. Then, T1w reference images were normalized to a study specific T1w template, before being normalized to the MNI template by rigid-body, affine, and non-linear registration.^39^ Using the obtained parameters and warp-field, the MTw and T1w-images were transformed to MNI-space. For details see Supplementary Methods.

### Structural signal extraction

As a measure of LC integrity, we calculated an MTw contrast-to-noise-ratio (CNR_MTw_).^16^ CNR_MTw_ maps were calculated from the normalized MTw images by subtracting the mean signal intensity (SI) of a reference region in the pontine tegmentum and dividing it by the reference region’s standard deviation:

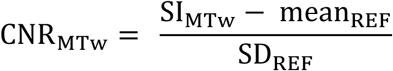

The LC region of interest (ROI) was the atlas from Ye et al.^37^ (5% probability), but excluding hyperintense voxels around the 4^th^ ventricle as described previously.^16^ In the present study, we further divided the LC mask across the middle along the rostro-caudal axis, forming four ROIs (right rostral, right caudal, left rostral, and left caudal) from which we extracted mean CNR_MTw_ (Figure 2B). This division was motivated in part by previous findings of predominantly caudal degeneration,^16,17^ but also to allow us to analyse the low signal-to-noise fMRI data at a sub-regional level while still being able to average across multiple voxels to increase sensitivity.

**Figure 2.**
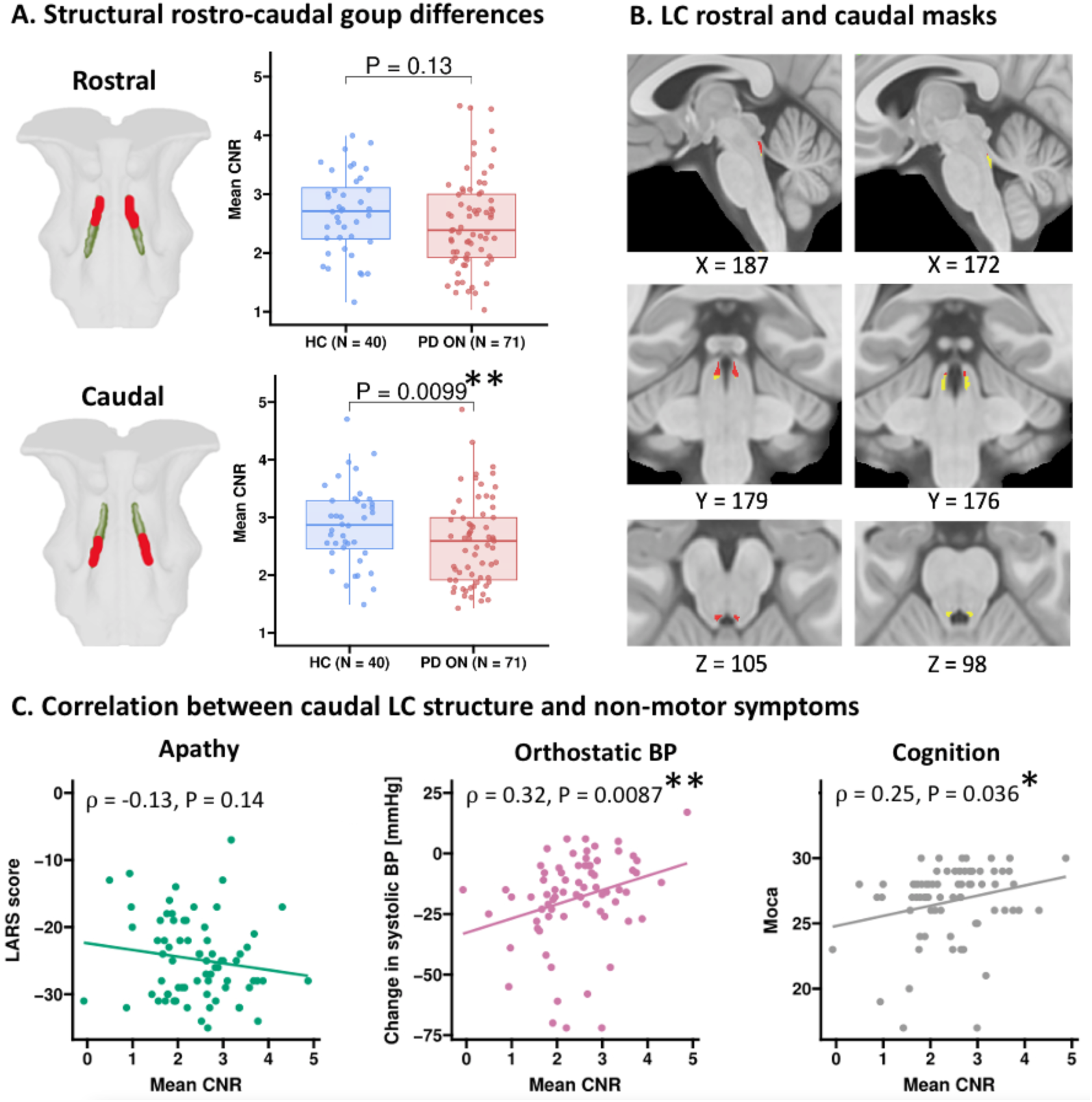
Rostro-caudal group differences for LC CNR_MTw_ and correlations to non-motor symptoms. (A) Group difference in rostral and caudal LC CNR_MTw_ for the structural group (N = 111). The p-values are derived from the mixed-effects regression model including CNR_MTw_ values in the LC for each side. For visualizing the effect, each boxplot shows one averaged data point per subject across sides. Compared to controls, patients had lower CNR_MTw_ in the caudal region (t_107.0_ = -2.63, P = 0.0099). There was no group difference in the rostral region (t_107.0_ = -1.54, P = 0.13). (B) Rostral and caudal LC masks for signal extraction. (C) Correlations between caudal structural LC integrity and non-motor symptoms for PD ON (N = 71). Structural integrity in the caudal LC region correlated positively with orthostatic changes in systolic BP and MoCA. There was no significant correlation with apathy. Abbreviations: CNR = contrast-to-noise ratio; LC = Locus Coeruleus; BOLD = blood-oxygen-level-dependent; PD ON = Parkinson’s patients on medication; PD OFF = Parkinson’s patients off medication; HC = healthy controls, BP = blood pressure; LARS = Lille Apathy Rating Scale; MoCA = Montreal Cognitive Assessment. *P < 0.05, **P < 0.01, ***P < 0.001.

### Functional image processing

Image processing of the fMRI data was performed using FMRIB Software Library (FSL 6.0.4), Statistical Parametric Mapping (SPM12), ITK-SNAP, and ANTs. First, the EPI slabs were distortion corrected using FSL-TOPUP. Second, we applied slice-time- and motion-correction with SPM. Next, we developed a co-registration pipeline based on *layerfMRI.com* (https://layerfmri.com/2019/02/11/high-quality-registration/) using ANTs. To achieve co-registration between low-resolution EPI and T1w images, we first normalized the skull-stripped T1w and the MTw images to an MNI template (rigid-body, affine, and non-linear registration). T1w images were then co-registered to EPI space with a rigid, affine, and non-linear registration, focusing the registration on the brainstem. Using the resulting transformation parameters, we then transformed our ROIs (four LC ROIs, a fourth ventricle ROI to derive CSF pulsations potentially affecting the LC BOLD signal and ROIs of the superior and inferior colliculi^40^) to individual EPI space. For details see Supplementary Methods.

### Functional signal extraction

We used SPM12 to create regressors and estimate their beta-weights. Regressors of interest were the onsets of arousal stimuli (noise and animals), time-binned into two regressors for the first 22 and last 23 stimuli to test for a potential habituation to the arousal stimuli over time. A third regressor modelled the onsets of target events (tones and flowers) where a response was registered. As nuisance regressors, we added respiration, pulse, head motion and signal from the 4^th^ ventricle. We used a high-pass filter (128 s) to remove slow signal drifts.

We used MARSeille Boîte À Région d’Intérêt (MarsBaR-0.45) to extract the average beta weights of the arousal stimuli regressors for each of the four LC ROIs and each participant,^41^ thus extracting signal from the same ROIs as in the structural MTw analysis.

### Statistical analyses

Statistical analyses were performed using R (version 4.2.2) using the mixed model package “lme4” for summary inference.^42^ Statistical significance was accepted at a threshold of p < 0.05. Residual plots and quantile-quantile plots were visually inspected and showed no apparent deviations from homoscedasticity or normality.

### Structural data from the extended cohort (n = 111)

#### Structural rostro-caudal pattern

For the structural MRI, group differences were tested using linear mixed models with CNR_MTw_ signal in the caudal and rostral LC as outcome variable, fixed effect factors ‘group’ (HC, PD) and ‘ROI laterality’ (right, left), ‘age’, ‘sex’ and random effect ‘subject’, with ‘group’ being the main variable of interest. It was defined in R syntax as follows:

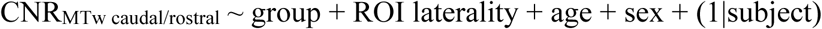

We also explored whether any group differences would be apparent across the whole LC, left and right LC separately, or within the left and right caudal sub-region separately with similar models (see Supplementary Methods).

#### Structural correlation to non-motor symptoms

To confirm our previous finding,^16^ we used the same method (one-tailed spearman analyses) to test correlations between structural MRI and non-motor symptoms in the extended cohort. Here, we hypothesised that caudal LC CNR_MTw_ would correlate negatively with apathy (LARS) and positively with orthostatic hypotension in PD.^16^ Further, we hypothesised that LC CNR_MTw_ would correlate positively with cognition (MoCA) based on multiple recent studies.^19–22^ Given the predominantly caudal degeneration established previously, we focused our analyses of the relationship between non-motor symptoms and structural LC changes on the caudal LC, with exploratory analyses of the rostral part. P-values were corrected for multiple comparisons with the Holm method.

### Functional data from the subgroup (n = 57)

#### Functional MRI quality check

To check the quality of the brainstem fMRI data, we conducted paired t-tests comparing activation in the inferior colliculus (receiving auditory inputs) and the superior colliculus (receiving visual inputs) during the auditory and visual arousal task.^43^ We averaged across the right and left colliculi masks and across sessions to increase robustness. Given the anatomical proximity of the colliculi to LC, this analysis served as a sanity check to confirm that the stimuli elicited the expected modality-specific neural responses. The analysis also offered a quality check for signal reliability in the surrounding brainstem region, including the LC. Additionally, we tested whether the expected response patterns (i.e., greater inferior colliculus activation by auditory stimuli and greater superior colliculus activation by visual stimuli) were present within each group. This ensured that any group differences in LC activation were not confounded by upstream group differences in sensory processing.

#### Functional rostro-caudal pattern

Similarly to the structural MRI, we used linear mixed models to test for rostral and caudal group differences in the functional response to arousal stimuli. The model was expanded, since the functional data was measured over two sessions with two paradigms. The outcome variable ‘functional LC response’ was the signal from each ROI extracted in MarsBaR. Fixed effect factors were ‘group’ (HC, PD ON, PD OFF), ‘session’ (first, second), ‘time-bin’ (first 22 and last 23 stimuli), ‘paradigm’ (auditory, visual) and ROI laterality (right, left). ‘Subject’ was used as random effect, and we controlled for age and sex:

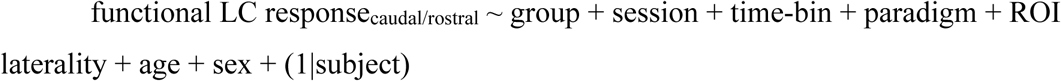

As for the structural data, we explored group differences across other ways of dividing the LC (see Supplementary Methods).

### Exploratory analysis

#### Structure-function relationship

To explore the relationship between structural disintegration of the LC and its functional activation, we used a cut-off of maximum six months (180 days) between the structural and functional scans to avoid major confounds of disease progression between the time points. Due to the pandemic and logistical and resource constraints, median duration between structural and functional data acquisition was 203 days (HC)/213 days (PD) (range: 9-636 days). Data had been acquired within 180 days for 10 HC and 12 PD patients who had performed both tasks.

Since structural MRI was only assessed in the ON-medication state, we first tested the structure-function relationship in the comparable state of dopaminergic integrity/restoration (HC and PD ON). We used regression models with the caudal and rostral LC response to arousal stimuli as dependent variables and the explanatory regressors ‘CNR_MTw_ signal’ and ‘group’ (HC, PD ON):

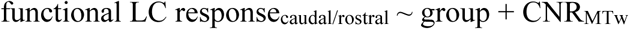

Since we had fewer subjects for this analysis, a model with a random effects subject factor could not be estimated. In order to use a regression model, we averaged the data to one structural and functional MRI data point per subject.

In the PD group, we then explored an interaction effect between dopaminergic state and LC structure on functional LC response:

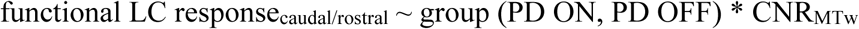

Finally, to explore the direction of above interaction, we correlated (two-sided spearman correlation) each patient’s LC structural integrity with the difference in functional LC response between ON and OFF state.

#### Functional correlation to non-motor symptoms

To examine the function-symptom relationship, we applied the same cut-off of six months between the clinical scoring and functional scans, providing the same sub-sample as above. Given the predominantly caudal degeneration, we focused our analyses on the caudal LC, with exploratory analyses of the rostral part. We used a regression model to test the effects of ‘group’ (HC, PD ON, PD OFF) and ‘functional LC response’ on non-motor symptoms:

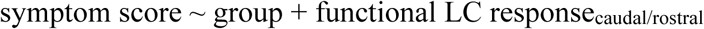

The test was run for each of the three symptoms (apathy, cognition, orthostatic hypotension). For orthostatic hypotension, the ‘group’ factor had only two levels as it was only measured in patients. ‘Functional LC response’ was the averaged signal across left and right side.

## Data availability

The pseudonymized data can only be shared with a formal Data Processing Agreement and a formal approval by the Danish Data Protection agency in line with the requirements of the General Data Protection Regulation.

## Results

### Clinical Data

There were no group differences in age, sex, MoCA or LARS scores (all *P-*values > 0.05, Table 1). Patients scored higher on the depression inventory than controls in the structural MRI group (n = 111, *P* = 0.00013) and fMRI subgroup (n = 57, 0.0099). In the structural MRI group, 29/71 patients had orthostatic hypotension defined as systolic BP drop ≥20 mmHg or a diastolic BP drop ≥10 mmHg.^44^ Ten patients met the criteria for both systolic and diastolic orthostatic hypotension, 16 for systolic hypotension only, and three for diastolic hypotension only. In the functional subgroup 8/30 patients had orthostatic hypotension with two patients meeting the criteria for both systolic and diastolic hypotension and six patients for systolic hypotension only.

### Structural Disintegration of Locus Coeruleus

#### Rostro-caudal pattern of disintegration

The caudal CNR_MTw_ signal was lower in PD ON compared to controls (t_107.0_ = -2.63, *P* = 0.0099), with no group difference in rostral LC (t_107.0_ = -1.54, *P* = 0.13) (Fig. 2A). In both rostral and caudal LC, there was a significant effect of laterality with both patients and controls showing higher signal on the right side (rostral: t_110.0_ = 7.96, *P* < 0.001; caudal: t_110.0_ = 8.32, *P* < 0.001; Supplementary Table 2).

Additional analyses (exploratory, no correction for multiple comparisons) of other divisions of the LC revealed lower structural signal in PD ON compared to controls across the whole (bilateral) LC (t_107.0_ = -2.25, *P* = 0.026), in the entire right LC (t_107.0_ = -2.82, *P* = 0.0057), and in right caudal LC (t = -3.08, *P* = 0.0026), but not for any other (sub-)regions (Supplementary Fig. 1& 2).

#### Correlations between LC disintegration and non-motor symptoms

Based on previous research, we expected a correlation between the structural mean CNR_MTw_ signal in the caudal LC region and three non-motor symptoms: apathy, orthostatic hypotension, and cognitive dysfunction.^16,19–22^ Mean CNR_MTw_ signal in the caudal LC correlated with orthostatic hypotension (rho = 0.32, Holm-corrected *P* = 0.0087) and MoCA scores (rho = 0.25, Holm-corrected *P* = 0.036; Fig. 2C). In contrast to our previous analysis, there was no significant correlation with apathy in this extended dataset (rho = -0.13, Holm-corrected *P* = 0.14). There were no significant correlations between rostral structural LC integrity and the three non-motor symptom clusters, nor between rostral or caudal structural LC integrity and depression or general non-motor symptoms (Supplementary Fig. 3).

### Functional activation evoked by arousing stimuli

#### Modality-specific activation of the superior and inferior colliculi

To test if the arousal stimuli elicited a reliable BOLD response in relevant brainstem regions, we tested the activation of the superior and inferior colliculi (Fig. 3). Auditory stimuli elicited higher activity in the inferior than superior colliculus (t_55_ = 5.56, *P* = 8.1^e-7^), while visual stimuli showed the reverse (t_54_ = 4.17, P = 0.00011). These differences were consistent within each group (Supplementary Fig. 4). There were no group differences in the superior colliculus to visual stimuli or in the inferior to auditory stimuli (Supplementary Fig. 5).

**Figure 3.**
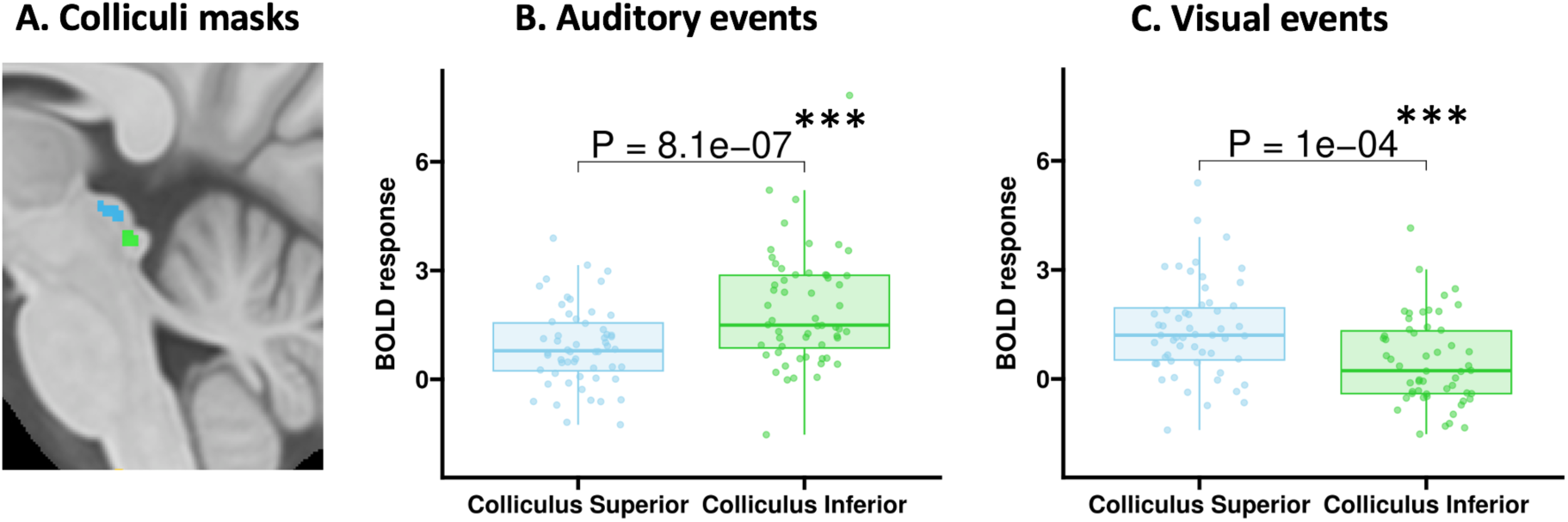
BOLD responses for the superior and inferior colliculi during auditory and visual stimuli. (A) Colliculi masks from Brainstem Navigator in MNI space: superior colliculus (blue) and inferior colliculus (green). (B) The inferior colliculus had significantly higher activity than the superior colliculus to auditory stimuli (paired t-test, t_55_ = 5.56, P = 8.1e^-7^). (C) The superior colliculus had significantly higher activity than the inferior colliculus to the visual stimuli (paired t-test, t_54_ = 4.17, P = 0.00011). Each boxplot shows one data point per subject averaged over sessions. Abbreviations: BOLD = blood-oxygen-level-dependent. *P < 0.05, **P < 0.01, ***P < 0.001.

#### Rostro-caudal activation pattern in locus coeruleus

As hypothesised, functional LC activation followed a rostro-caudal gradient matching the structural pattern (Fig. 4). Specifically, BOLD responses in the caudal LC were significantly lower in the PD ON-medication group compared to controls (t_71.6_ = -2.58, *P* = 0.012). Although caudal LC responses were also reduced in PD patients in the OFF-medication state, this difference did not reach significance (t_73.5_ = -1.64, *P* = 0.10). Consistent with structural findings, no significant group differences were observed in rostral LC activation.

**Figure 4.**
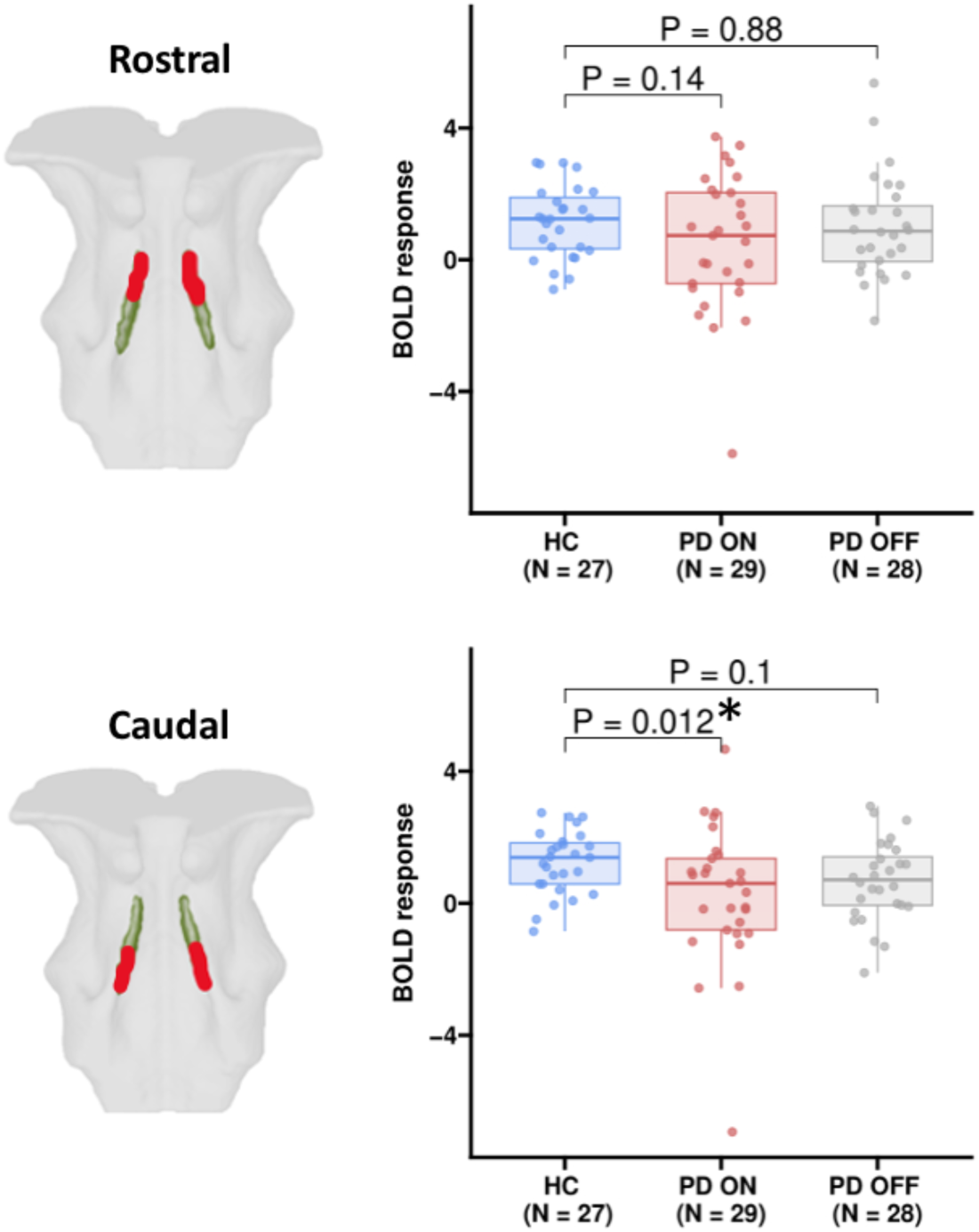
Rostro-caudal group differences for LC fMRI. Group difference in rostral and caudal LC BOLD response for the fMRI subgroup (N = 57). The p-values are derived from a mixed-effects regression model including contrast values in the LC for each side, each session, each time bin and each task (see Methods). For visualizing the effect, each boxplot shows one averaged data point per subject across side, time bin and task (and across sessions for HC). In the caudal region, PD ON showed reduced BOLD response compared to controls (t_71.6_ = -2.58, *P* = 0.012). The group difference between PD OFF and controls was not significant (t_73.5_ = -1.64, *P* = 0.10). There was no group difference for the rostral region. Abbreviations: LC = Locus Coeruleus; BOLD = blood-oxygen-level-dependent; PD ON = Parkinson’s patients on medication; PD OFF = Parkinson’s patients off medication; HC = healthy controls. *P < 0.05, **P < 0.01, ***P < 0.001.

For both structural and functional MRI, exploratory analyses revealed significant differences between healthy controls and patients in the ON-medication state, when considering the average signal in the right LC and throughout the entire LC (Supplementary Fig. 1). BOLD responses to arousing stimuli were significantly reduced in the PD ON-medication group compared to controls across the entire LC (t_59.4_ = -2.42, *P* = 0.019) and in right LC (t_66.2_ = -2.33, *P* = 0.023), but not in left LC (t_71.4_ = -1.85, *P* = 0.068). Within the caudal LC, right-sided responses were significantly lower in patients in the ON-medication state (t_81.6_ = -2.42, *P* = 0.018; Supplementary Fig. 2).

A significant main effect of ‘time-bin’ indicated habituation to arousal stimuli in the caudal LC, with reduced BOLD responses during the second half of trials compared to the first half (t_757.8_ = -3.49, *P* < 0.001; Supplementary Table 2). Additionally, we found a significant main effects of ‘paradigm’. Visual stimuli evoked lower BOLD responses than auditory stimuli in both rostral (t_774.4_ = -2.09, *P* = 0.037) and caudal LC (t_777.6_ = -2.50, *P* = 0.012).

#### Structure-function relationship

We examined the structure-function relationship in a small subgroup of participants (healthy controls: n = 10; patients: n = 12) for whom structural and functional brain scans had been acquired within six months. In this subgroup, structural and functional signals were positively correlated in caudal LC (t = 3.03, *P* = 0.0069; Fig. 5A). No significant structure– function relationship was observed in rostral LC (Supplementary Fig. 6A).

**Figure 5.**
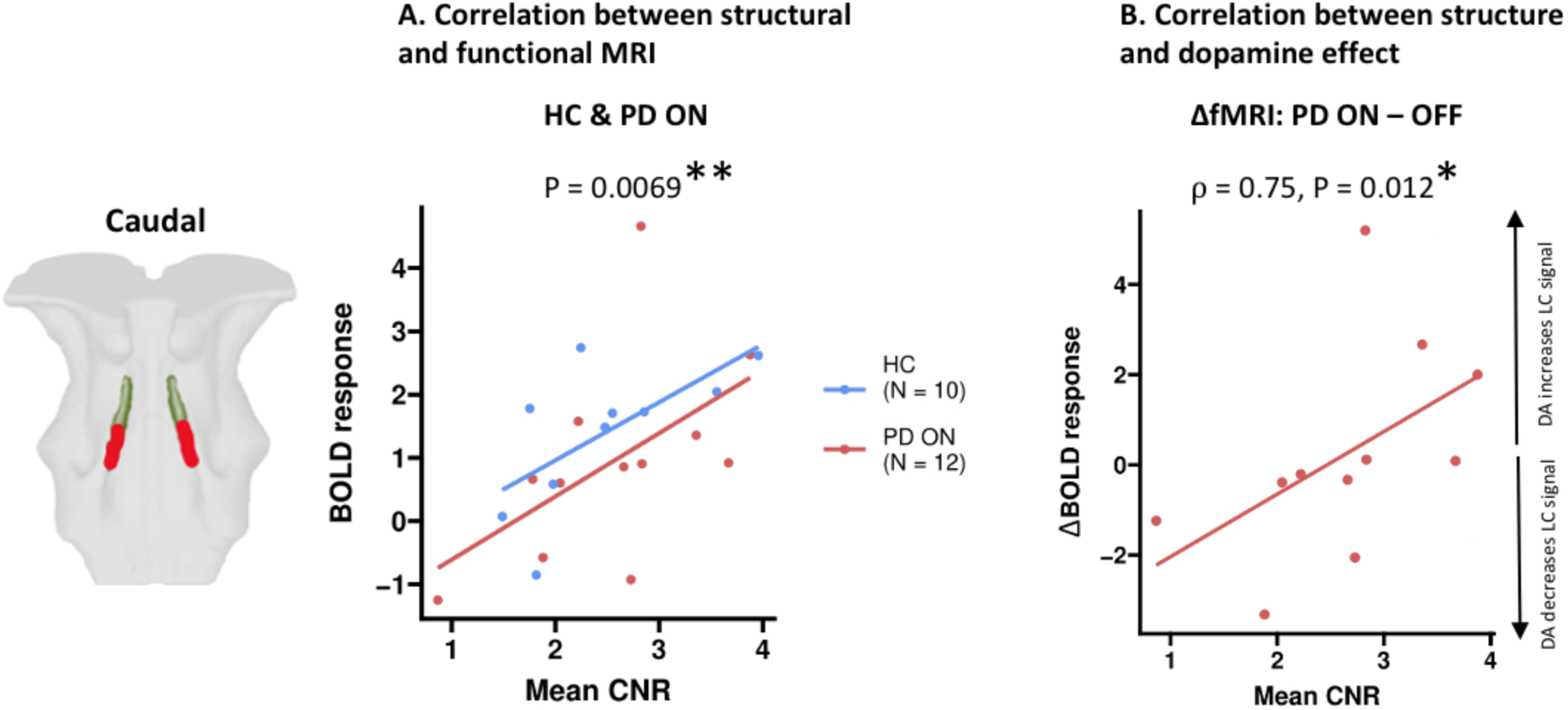
Exploration of structure-function relationship and dopamine effect. BOLD response is averaged to one data point per subject. (A) Scatterplot showing the positive correlation between structural and functional signal with best-fitting regression line for the data points. Note that the p-value refers to the regression coefficient from the linear model also containing a group factor (see text). The analysis was performed on subjects in their habitual state, meaning patients on their normal dopaminergic medication and controls with natural dopamine in the brain. (B) Scatterplot showing the significant correlation between medication-induced change in functional LC response (*on*-state minus the *off*-state) and structural LC integrity with best-fitting regression line for the data points (rho = 0.75, P = 0.012). The correlation suggests that in patients with lower structural LC integrity, dopaminergic medication decreases LC activity while it has the opposite effect in patients with relatively high LC integrity. Abbreviations: (LC: locus coeruleus). *P < 0.05, **P < 0.01, ***P < 0.001.

In patients, withdrawal of dopaminergic medication significantly altered the structure–function relationship, reflected in a significant interaction effect between dopaminergic state and structural integrity on LC function (t = 2.17, *P* = 0.042). Notably, the effect of medication on LC activation (ON-medication minus OFF-medication state) varied as a function of structural LC integrity: Patients with lower neuromelanin signal in caudal LC (i.e., lower structural integrity) showed reduced functional activation with dopaminergic medication, whereas those with higher caudal neuromelanin signal (i.e., higher structural integrity) showed increased activation (rho = 0.75, *P* = 0.012; Fig. 5B). These effects were not present in the rostral LC (Supplementary Fig. 6B & C).

#### Correlations to non-motor symptoms

We explored the relation between LC function and non-motor symptoms in the subset of participants, where functional scans were obtained within six months after clinical assessment. As this is the first study to examine subregional LC activity in relation to non-motor symptoms, both rostral and caudal LC responses were analysed. Rostral but not caudal LC responses to arousing stimuli were associated with the degree of apathy (t = 2.99, *P* = 0.0055) with no group difference. Neither MoCA scores nor orthostatic hypotension showed significant associations with functional LC responses to arousing stimuli.

## Discussion

This multimodal ultra-high field MRI study yields three key insights into LC degeneration and dysfunction in PD: First, using event-related fMRI, we provide the first evidence of impaired LC reactivity to arousal stimuli in PD. Reactivity was impaired in the caudal part of the LC and reached significance in the ON-medication state. Second, using neuromelanin-sensitive structural MRI, we confirmed a selective caudal loss of neuromelanin signal in the LC in a larger sample. We further found a positive correlation between the structural and functional signals in caudal LC in a subgroup of medicated PD patients and healthy controls. Within the PD group, the dopaminergic medication state modulated this structure-function relationship. Third, structural and functional LC signals were linked to non-motor symptoms in PD, highlighting the clinical relevance of noradrenergic dysfunction beyond dopaminergic pathology.

### Rostro-caudal functional impairment of locus coeruleus

Noradrenergic neurons in the LC increase firing in response to salient stimuli and play a crucial role in arousal.^4,45–49^ Their degeneration is therefore expected to reduce BOLD responses to arousing sensory input. Consistent with this hypothesis, patients displayed a significantly reduced arousal response in the caudal LC in the ON-medication state. A similar but non-significant trend was found in the OFF-medication state. To date, only two studies with possibly overlapping samples have reported altered LC resting-state connectivity in PD.^18,50^ However, this is the first event-related fMRI study to demonstrate impaired LC activation in response to arousal stimuli in PD.

The functional impairment was localized in the caudal LC, mirroring the spatial pattern of structural disintegration as revealed by structural MRI. The absence of group differences in the responses of the superior and inferior colliculi to visual and auditory stimuli (Fig. 4, Supplementary Fig. 4 & 5) indicates that the observed difference in LC arousal response is not due to an upstream gating problem.

At the subregional level, an fMRI study in healthy adults has investigated LC activation, showing an increased response in the rostral LC to auditory oddball stimuli.^51^ Since the rostral LC is densely connected to associative cortical areas in humans and animals, ^52–56^ the rostral activation of LC may reflect a “difference” signal that is conveyed from LC to associative cortical areas. In our data, both caudal and rostral LC showed significant arousal-related activation across groups, but no significant rostro-caudal differences, possibly because the stimuli did not signal any deviation from the ongoing flow of sensory input as in an oddball setting. It may thus be expected that arousing stimuli may lead to differential spatial engagement of the LC depending on the context.

### Rostro-caudal pattern of structural degeneration

Extending the sample size, we corroborated our previous finding showing a predominantly caudal degeneration in PD. Tracing studies have shown that the LC is not uniformly organized. Its rostro-dorsal part projects primarily to neocortical and medial temporal regions,^55^ while its caudo-ventral parts project primarily to the cerebellum and spinal cord.^56^ Degeneration also follows a spatial gradient across diseases: In Alzheimer’s disease^57,58^ and healthy aging,^59^ LC degeneration occurs predominantly in the rostral portion of LC, whereas in PD, histological^60^ and imaging studies^16,17,19,36,61^ show converging evidence that neurodegeneration occurs predominantly in the caudal LC. Our results confirm this caudal vulnerability of LC in PD. This selective degeneration may reflect pathological processes such as alpha-synuclein transmission via vagal projections from the gastrointestinal system^62^ or increased exposure to environmental toxins near the fourth ventricle.^1,19^ Of note, the right-lateralized loss of neuromelanin contrast observed in our sample may be attributable to scanner-specific field inhomogeneities, as previously reported.^63,64^

We explored the observed overlap of structural and functional alterations in caudal LC in a subgroup of participants. Due to the progressive nature of the disease, we limited this analysis to data acquired within six months and to data from subjects’ habitual state, i.e. patients ON dopaminergic medication and controls with natural levels of dopamine. In medicated patients and healthy controls, the caudal but not the rostral LC showed significant correlations between the regional neuromelanin signal and the functional response to arousing stimuli (Fig. 5, Supplementary Fig. 6A). To our knowledge, only one other MRI study has investigated the structure-function relationship in the LC in healthy adults only and found no significant associations.^51^ This may be due to differences in task design or the inclusion of a predominantly young sample in whom neuromelanin accumulation is still limited.^6,7^ In contrast, our study provides the first evidence of a structure-function correlation in the human LC. However, given the small sample size of this exploratory analysis, these findings should be interpreted with caution and require independent replication.

#### Impact of dopaminergic medication on noradrenergic arousal function

Since levodopa is a precursor of both noradrenaline and dopamine, its administration might thus increase noradrenaline levels in the brain. Accordingly, dopaminergic medication affects many non-motor systems in PD commonly associated with noradrenaline.^65,66^ Animal studies testing the effect of levodopa on noradrenaline levels are inconclusive,^67^ with some reporting increases,^68,69^ but others no effects or even decreases.^70–72^ In parkinsonian animal models, levodopa tends to decrease noradrenaline levels in different brain regions.^69,73^ Thus, both dopamine depletion and medication may affect the noradrenergic system. These interactions are relevant to the interpretation of our findings. Notably, we found a differential effect of medication on LC function in a small subgroup of patients that depended on the degree of LC degeneration during PD (Fig 5B). Dopaminergic medication increased the caudal LC response to arousing stimuli in patients with a structurally more intact LC. Conversely, levodopa decreased LC activation, in patients with stronger degeneration. This pattern aligns with preclinical evidence of levodopa-induced noradrenaline reduction in PD animal models.^69,73^ However, this analysis is based on a small sub-sample, limiting its robustness and requiring independent replication.

#### Associations between alterations of locus coeruleus and non-motor symptoms

In our extended cohort, we confirmed the previously reported association between caudal LC degeneration and orthostatic hypotension^16^ and additionally identified a significant correlation with cognitive impairment. Orthostatic hypotension is a common autonomic dysfunction in PD, affecting 30-58% of people with PD^74^ (40.8% in our sample) and may result from both central or peripheral noradrenergic dysfunction.^75^ Several findings support a central origin of orthostatic hypotension in PD, including associations between noradrenaline transporter density in the LC and orthostatic hypotension,^76^ reduced LC-mediated inhibition of parasympathetic cardiac innervation,^77–81^ and co-occurring degeneration of peripheral postganglionic sympathetic neurons and the LC.^82,83^ The specific involvement of the caudal LC is anatomically plausible, as this region projects to the spinal cord^56,84,85^ where it modulates sympathetic preganglionic neurons.^86^ A recent MRI study in older healthy adults also linked neuromelanin signal in caudal LC to cardiac responsivity to acute stress.^87^ Together with our finding, these results support the role of caudal LC in peripheral autonomic regulation and underscores the importance of sub-regional assessments with MRI.

We also found that a reduced neuromelanin signal in caudal LC correlated with lower MoCA scores, aligning with previous studies that have linked reduced LC integrity with cognitive impairment in PD.^19–22^ Cognitive deficits are already present in around 40% of PD patients at the time of clinical diagnosis.^88^ Apart from fronto-executive deficits, cognitive deficits such as working memory respond poorly to dopamine therapy.^89^ This has been attributed to neurodegeneration of other transmitter systems, including LC-mediated noradrenergic signaling. ^20,22^

As several previous reports,^90–93^ the severity of cognitive impairment correlated positively with the severity of orthostatic hypotension in the PD group (rho = 0.33, *P* = 0.0026) (Supplementary Fig. 8). Two hypotheses have been proposed to explain this relationship,^75^ (1) a vascular hypothesis where cerebral hypoperfusion leads to neural damage, and (2) a shared underlying neuroanatomical or neurochemical basis such as LC degeneration. Our findings, showing that both symptoms correlate with caudal LC degeneration, support the latter explanation and point to a common noradrenergic mechanism.

Extending previous findings that linked apathy to LC degeneration,^16,19,94^ the BOLD responses in rostral LC were significantly associated with apathy across all participants who underwent ultra-high field fMRI (Fig. 6). Neuroanatomically, apathy has been suggested to be related to prefrontal-basal ganglia network dysfunction,^95–97^ regions that are predominantly innervated by the rostral LC.^56,84,98^ Interestingly, we found that higher rostral activity correlated with a higher apathy score, indicating an association with hyperarousal rather than reduced LC activation. However, as this is an exploratory finding based on a small sample, replication in larger cohorts is needed to confirm and clarify the directionality and clinical implications of this association.

**Figure 6.**
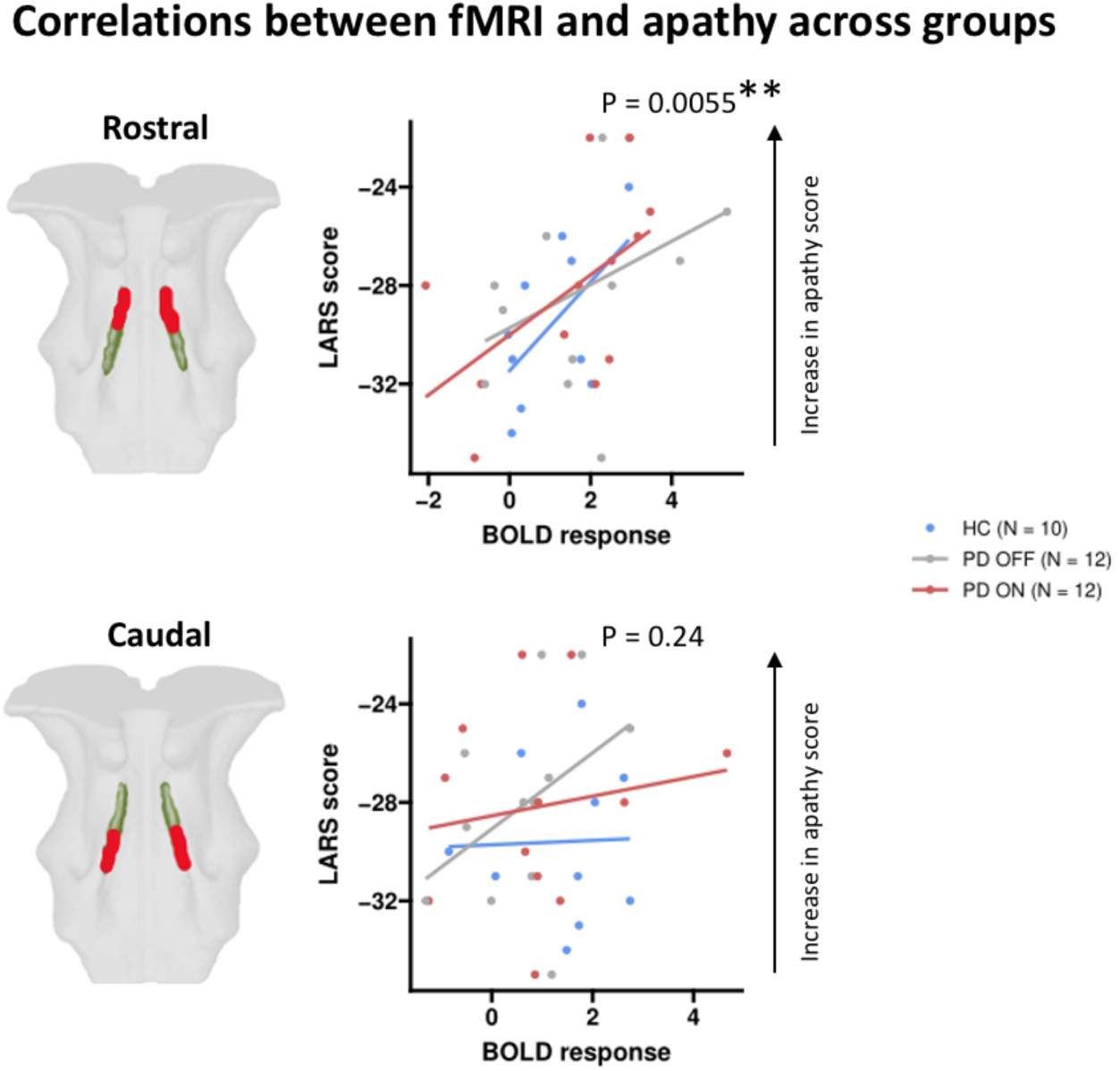
Association between rostral and caudal LC function and apathy. Scatterplots showing the correlations between apathy scores and functional LC response (averaged to one data point per subject) in the rostral (top row) and caudal (bottom) LC for all three groups with best-fitting regression lines for the data points. Note that the p-values refer to the regression coefficient for the ‘functional LC response’ factor from the linear models explaining the apathy score (see text). Rostral LC activity was significantly associated with apathy across groups (P = 0.0055). There was no effect of group and no significant association with caudal LC activity (all P-values > 0.05). As for the structure-function correlation, we used a cut-off of maximally six months between the assessment of the clinical scores and functional scans. *P < 0.05, **P < 0.01, ***P < 0.001.

### Strengths and limitations

Leveraging the enhanced signal-to-noise ratio of ultra-high field MRI enabled us to investigate disease-induced structural and functional LC alterations at the subregional level. Rigorous co-registration and correction procedures, including adjustments for physiological confounds such as CSF signal, cardiac pulsations, and respiratory motion, combined with signal validation in other brain-stem nuclei, allowed us to measure the functional arousal response in this small and anatomically challenging region.^100^ Using two different arousal paradigms with auditory and visual arousal stimuli provided internal validation regarding feasibility of ultra-high field brainstem fMRI as evidenced by modality-specific activation patterns in the inferior and superior colliculi. The unpredictability of stimulus presentation increased salience, and the absence of group differences in collicular responses supports the conclusion that LC differences were not due to perceptual deficits.

However, several limitations must be acknowledged. First, the right-left asymmetry in the LC MRI signal might reflect the field inhomogeneity previously reported in Philips scanners rather than an actual neurobiological difference.^63,64^ Second, while the MTw signal has been closely linked to neuromelanin-containing neurons in prior studies,^101–103^ the MTw signal is sensitive, but not necessarily specific to neuromelanin. It may also reflect other tissue properties such as water content or macromolecular composition. Thus, reduced signal reflects reduced LC integrity, but not exclusively a selective loss of neuromelanin. Thus, a reduced MTw signal reflects reduced LC integrity, but not exclusively a regional loss of neuromelanin.^15,103,104^ Third, the variable time interval between acquisition of structural and functional data reduced the number of participants eligible for structure–function correlation analyses, thereby limiting statistical power. Fourth, the OFF-medication state in PD patients was pragmatic and does not represent a complete withdrawal. Residual longer lasting dopaminergic effects cannot be ruled out.^105^ Furthermore, antihypertensive medication was not stopped prior to orthostatic blood pressure measurements, which may have influenced the interpretation of autonomic findings. Lastly, while the MoCA is a widely used cognitive screening tool, it may lack sensitivity for detecting early mild cognitive impairment in PD.^22^

### Conclusion

PD-related degeneration of the locus coeruleus follows a rostro-caudal gradient with the caudal LC being most affected. LC reactivity to arousing stimuli also is reduced in caudal LC in PD. Importantly, both structural and functional LC measures were linked to non-motor symptoms in PD, underscoring the contribution of noradrenergic dysfunction to clinical impairment beyond dopaminergic pathology.

## Acknowledgements

We thank Associate Professor Esben T. Petersen and Vincent O. Boer for their support in developing the MRI acquisition protocols, Søren A. Fuglsang for assistance with experimental design, data processing, and analysis, and James B. Rowe for his valuable manuscript feedback. Lastly, we would like to thank all the patients and healthy controls for their participation in this project.

## Supplementary material

Supplementary material is available online.

