## Supplementary Material for "Blunted response of caudal locus coeruleus to arousing stimuli in Parkinson’s Disease"

### SUPPLEMENTARY METHODS:

#### Study Procedure

Details on audiogram for eligibility for fMRI sessions:

To assess eligibility for the functional sessions, a pure-tone audiogram was performed with an Oscilla USB 31 audiometer, testing hearing thresholds at the following carrier frequencies: at 100 Hz, 250 Hz, 500 Hz, 1000 Hz, 2000 Hz, 4000 Hz and 8000 Hz.

#### Event-related arousal paradigms

Details on auditory stimuli in auditory paradigm:

The auditory paradigm (played with Matlab R2017a) consisted of 45 bursts of low-pass filtered Gaussian white noise at 6 kHz using a 2nd-order Butterworth filter. Each burst had a duration of 500 ms and a sound pressure level (SPL) of 95 dB. Inter-stimulus-intervals were always greater than 10 s to allow BOLD responses to peak before the next stimulus and no stimuli were presented during the first 20 s. Target tones were played for 500 ms with carrier frequencies of 1 kHz and SPL of 85 dB. Target tones were never played as one of the first 4 stimuli or the last stimulus.

#### Magnetic Resonance Imaging Data

Details on structural images acquired on experimental day 2:

First, to guide co-registration and normalization, we acquired T1-weighted (T1w) Magnetization Prepared RApid Gradient Echo (MPRAGE) images with a field-of-view (FOV) covering the entire head (voxel size: 1 mm isotropic, number of voxels: 200 x 288 x 288, echo time = 2.2 ms, repetition time = 4.9 ms, acquisition time = 3:30 minutes). Second, as a measure of LC integrity, we acquired magnetization transfer-weighted (MTw) images using a 3D high-resolution, ultra-fast gradient echo sequence aligned to the AC-PC line with a FOV covering the midbrain and rostral pons (voxel size 0.4x0.4x1.0 mm, number of voxels: 640 x 640 x 34, echo time = 4.1 ms, repetition time = 8.1 ms, flip angle = 7 degrees, 2 averages). To achieve MT-saturation we applied 16 block-shaped pre-pulses (frequency offset: 2 kHz, flip angle = 278 degrees, duration = 10 ms, acquisition time = 8 minutes).

**Structural Image Processing**

Details on processing of MRI data acquired on experimental day 2:

The processing was performed with Advanced Normalization Tools (ANTs) software (http://stnava.github.io/ANTs/). MT-weighted and T1-weighted images were N4 bias field corrected for low-frequency spatial intensity inhomogeneities (5 resolution levels, number of iterations per level: 50 x 50 x 30 x 20, convergence threshold: 1 x 10-6, isotropic sizing for b-spline fitting: 200).[1] Within-subject co-registration and normalization to template space was performed by rigid-body, affine, and non-linear registration. First, to evaluate registration accuracy, high resolution images were transformed to individual subjects' T1w space. Then, T1w reference images were normalized to a study specific T1w template, before being normalized to the MNI 0.5 mm ICBM152 T1w asymmetric template by rigid-body, affine, and non-linear registration.33 Using the obtained co-registration and normalization parameters and warp-field, the high resolution MTw and T1w-images were transformed to MNI-space.

**Functional Image Processing**

Details on processing of fMRI data acquired on experimental days 3 and 4:

Image processing of the fMRI data was performed using FMRIB Software Library (FSL 6.0.4), Statistical Parametric Mapping (SPM12), ITK-SNAP, and ANTs .

First, the EPI slabs were distortion corrected using FSL-TOPUP. To estimate the distortion, a displacement field was calculated from the reversed phase encoding images and applied to the EPIs. Second, in SPM the EPI slabs were slice-time corrected and realigned to the first image, and the T1w reference images from the same session were skull-stripped.

Next, we developed a co-registration pipeline based in part on the steps described in layer fMRI blog (https://layerfmri.com/2019/02/11/high-quality-registration/). Here, ROIs defined in MNI space were coregistered to the individual EPI space instead of normalizing EPI images to MNI space, thus avoiding changing the EPI slabs due to non-linear registrations. To do this, we normalized the skull-stripped T1w images to the same MNI template as the MTw data using a rigid-body, affine, and non-linear registration. We then defined a brainstem mask in MNI space using ITK-SNAP and warped it to the individual T1w space. Using the brainstem mask to optimize the registration in our area of interest, T1w images were co-registered to EPI space with a rigid, affine, and non-linear registration. Having the transformation parameters from MNI to T1w and from T1w to EPI, we transformed the four LC ROIs from MNI to EPI space. Further, we defined and transformed a region inside the fourth ventricle (to derive potential CSF pulsations affecting the LC BOLD signal) as well as masks of colliculi inferior and superior (thresholded binary 0.35) from BrainStem Navigator (https://www.nitrc.org/projects/brainstemnavig/).[2]

If there were any misalignments in the co-registration, ITK-SNAP was used to manually change the initial alignment. The accuracy of the co-registration was checked individually for all images by AEL.
We used unsmoothed images to avoid smoothing signal from the bordering 4th ventricle into the LC.

**Statistical analyses of structural MRI data**

The two main models for testing group differences in the structural MRI data were:

$CNR_{MTw\ caudal}$ ~ group + ROI laterality + age + sex + (1|subject)

$CNR_{MTw\ rostral}$ ~ group + ROI laterality + age + sex + (1|subject)

Subsequently, we explored group effects in other divisions of the LC ROI using the following models:

$CNR_{MTw\ whole\ LC}$ ~ group + ROI + age + sex + (1|subject)

$CNR_{MTw\ left}$ ~ group + ROI + age + sex + (1|subject)

$CNR_{MTw\ right}$ ~ group + ROI + age + sex + (1|subject)

$CNR_{MTw\ left\ caudal}$ ~ group + age + sex + (1|subject)

$CNR_{MTw\ right\ caudal}$ ~ group + age + sex + (1|subject)

**Statistical analyses of functional MRI data**

The two main models for testing group differences in the functional MRI data were:

functional LC $response_{caudal}$ ~ group + session + time-bin + paradigm + ROI laterality + age + sex + (1|subject)

functional LC $response_{rostral}$ ~ group + session + time-bin + paradigm + ROI laterality + age + sex + (1|subject)

Subsequently, we explored group effects in other divisions of the LC ROI using the following models:

functional LC $response_{whole}$ ~ group + session + time-bin + paradigm + ROI + age + sex + (1|subject)

functional LC $response_{left}$ ~ group + session + time-bin + paradigm + ROI + age + sex + (1|subject)

functional LC $response_{right}$ ~ group + session + time-bin + paradigm + ROI + age + sex + (1|subject)

functional LC $response_{left\ caudal}$ ~ group + session + time-bin + paradigm + age + sex + (1|subject)

functional LC $response_{right\ caudal}$ ~ group + session + time-bin + paradigm + age + sex + (1|subject)

### REFERENCES SUPPLEMENTARY METHODS:


1. N. J. Tustison, B. B. Avants, P. A. Cook, et al. N4ITK: Improved N3 Bias Correction. *IEEE Transactions on Medical Imaging*. 2010;29(6):1310-1320. doi:10.1109/TMI.2010.2046908

2. García-Gomar MG, Bianciardi M. In vivo Probabilistic Structural Atlas of the Inferior and Superior Colliculi, Medial and Lateral Geniculate Nuclei and Superior Olivary Complex in Humans Based on 7 Tesla MRI. *Frontiers in Neuroscience*. 2019;13.


### SUPPLEMENTARY TABLES:

#### Supplementary Table 1

| | **Auditive Day 3** | **Auditive Day 4** | **Visual Day 3** | **Visual Day 4** |
|---|---|---|---|---|
| HC | 24 | 22 | 26 | 24 |
| PD ON (OFF) | 14 (14) | 15 (14) | 13 (13) | 14 (13) |
| **Total** | **52** | **51** | **52** | **51** |

**Supplementary table 1** Table of data per fMRI task per session

#### Supplementary Table 2

| | **Mixed model for the rostral LC (structural)** | | |
|---|---|---|---|
| *Predictors* | *Estimates* | *CI* | *p* |
| (Intercept) | 1.65 | 0.76 – 2.54 | **<0.001** |
| Group | -0.24 | -0.54 – 0.07 | 0.13 |
| ROI laterality | 0.47 | 0.35 – 0.59 | **<0.001** |
| Age | 0.01 | -0.00 – 0.02 | 0.086 |
| Sex | 0.11 | -0.18 – 0.41 | 0.45 |
| **Random Effects** | | | |
| $\sigma^2$ | 0.19 | | |
| $\tau_{00\ \text{participant\_id}}$ | 0.48 | | |
| ICC | 0.71 | | |
| $N_{\text{participant\_id}}$ | 111 | | |

| | |
|---|---|
| Observations | 222 |
| Marginal $R^2$ / Conditional $R^2$ | 0.107 / 0.745 |

| | Mixed model for the caudal LC (structural) | | |
|---|---|---|---|
| *Predictors* | *Estimates* | *CI* | *p* |
| (Intercept) | 2.96 | 2.00 – 3.92 | **<0.001** |
| Group | -0.44 | -0.76 – -0.11 | **0.0099** |
| ROI laterality | 0.60 | 0.46 – 0.74 | **<0.001** |
| Age | -0.00 | -0.02 – 0.01 | 0.64 |
| Sex | -0.30 | -0.62 – 0.02 | 0.070 |
| **Random Effects** | | | |
| $\sigma^2$ | 0.29 | | |
| $\tau_{00\ participant\_id}$ | 0.53 | | |
| ICC | 0.65 | | |
| $N_{participant\_id}$ | 111 | | |
| Observations | 222 | | |
| Marginal $R^2$ / Conditional $R^2$ | 0.161 / 0.706 | | |

**Supplementary table 2** Table of mixed models for the structural rostro-caudal analysis shown in Fig. 2.

**Supplementary Table 3**

| | Mixed model for the rostral LC (functional) | | |
|---|---|---|---|
| *Predictors* | *Estimates* | *CI* | *p* |
| (Intercept) | 2.54 | 0.31 – 4.77 | **0.027** |
| PD OFF | -0.06 | -0.85 – 0.74 | 0.88 |
| PD ON | -0.60 | -1.39 – 0.20 | 0.14 |
| Session | -0.18 | -0.60 – 0.24 | 0.39 |
| Time bin | -0.00 | -0.41 – 0.41 | 0.99 |
| Paradigm | -0.44 | -0.85 – -0.03 | **0.037** |
| ROI laterality | -0.24 | -0.65 – 0.17 | 0.26 |
| Sex | -0.52 | -1.29 – 0.25 | 0.18 |

| | | | |
|---|---|---|---|
| Age | -0.01 | -0.05 – 0.02 | 0.49 |
| **Random Effects** | | | |
| $\sigma^2$ | 8.98 | | |
| $\tau_{00}$ Subject_ID | 1.29 | | |
| ICC | 0.13 | | |
| N Subject_ID | 57 | | |
| Observations | 824 | | |
| Marginal $R^2$ / Conditional $R^2$ | 0.021 / 0.144 | | |

| | **Mixed model for the caudal LC (functional)** | | |
|---|---|---|---|
| *Predictors* | *Estimates* | *CI* | *p* |
| (Intercept) | 2.39 | 0.54 – 4.24 | **0.012** |
| PD OFF | -0.56 | -1.24 – 0.12 | 0.10 |
| PD ON | -0.87 | -1.55 – -0.20 | **0.012** |
| Session | -0.05 | -0.49 – 0.38 | 0.81 |
| Time bin | -0.77 | -1.20 – -0.34 | **0.001** |
| Paradigm | -0.56 | -0.99 – -0.12 | **0.012** |
| ROI laterality | 0.16 | -0.28 – 0.59 | 0.48 |
| Sex | -0.15 | -0.78 – 0.48 | 0.64 |
| Age | -0.01 | -0.04 – 0.02 | 0.59 |
| **Random Effects** | | | |
| $\sigma^2$ | 10.05 | | |
| $\tau_{00}$ Subject_ID | 0.59 | | |
| ICC | 0.06 | | |
| N Subject_ID | 57 | | |

| | |
|---|---|
| Observations | 824 |
| Marginal $R^2$ / Conditional $R^2$ | 0.035 / 0.088 |

**Supplementary table 3** Table of mixed models for the functional rostro-caudal analysis shown in Fig. 4.

### SUPPLEMENTARY FIGURES:

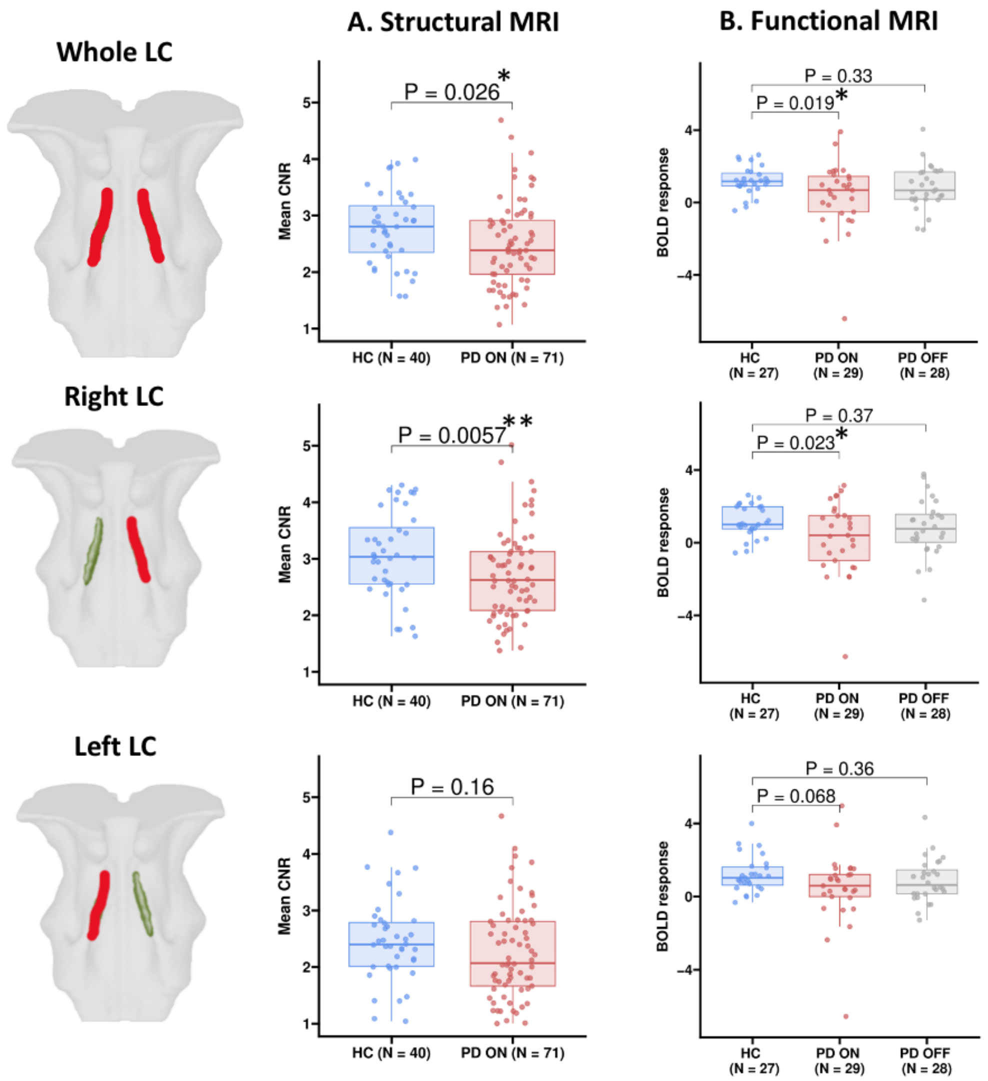


**Supplementary figure 1.** Similar to figure 2A, but for the whole, left, and right LC. It shows a significant group difference between HC and PD ON for the right LC and across the whole LC for both structural and functional MRI. The left side group difference is not significant on any of the modalities. *P < 0.05, **P < 0.01, ***P < 0.001.

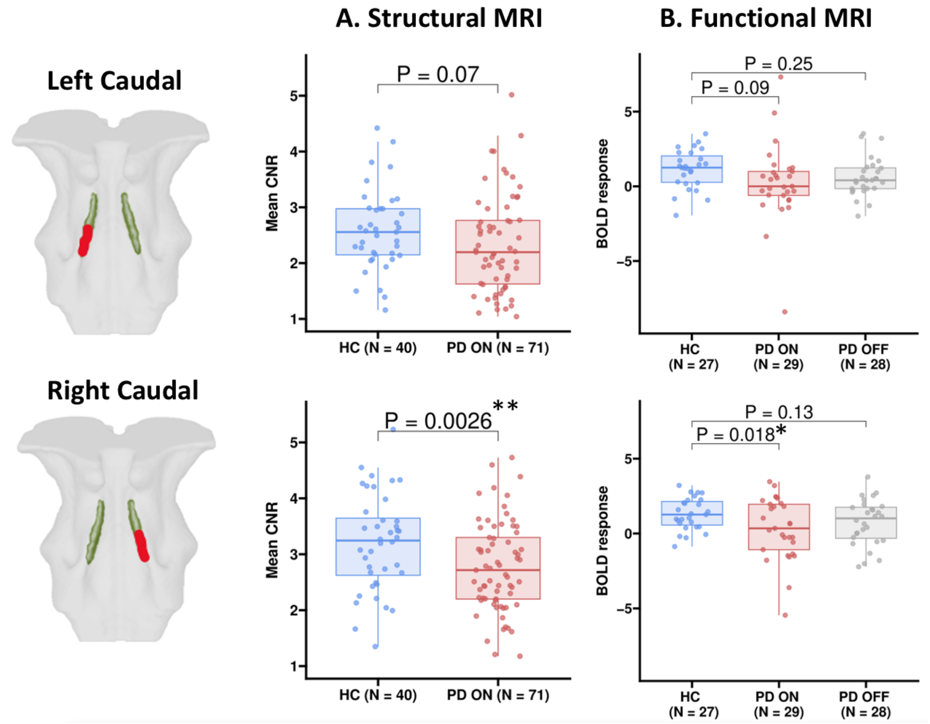


**Supplementary figure 2.** Similar to figure 2A, but exploring group differences within the caudal region (left and right). It shows a significant group difference between HC and PD ON for the right caudal LC for both structural and functional MRI. The left caudal group difference is not significant on any of the modalities. *P < 0.05, **P < 0.01, ***P < 0.001.

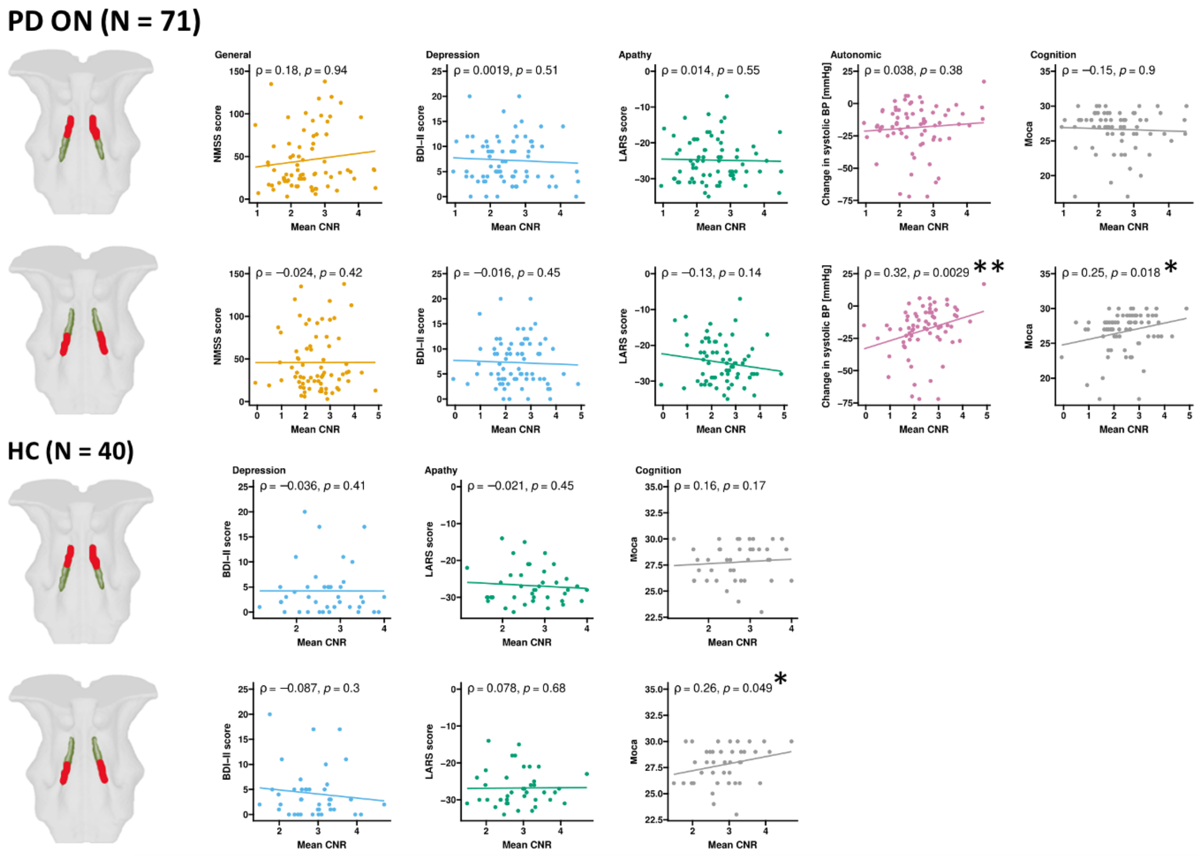

**Supplementary figure 3. Extended correlations between Mean CNR and symptoms for the extended cohort.** Similar to figure 2C, but includes rostral regions, correlations to NMSS and depression, and correlations for healthy controls. Unlike figure 2C, P-values are uncorrected. *P < 0.05, **P < 0.01, ***P < 0.001.

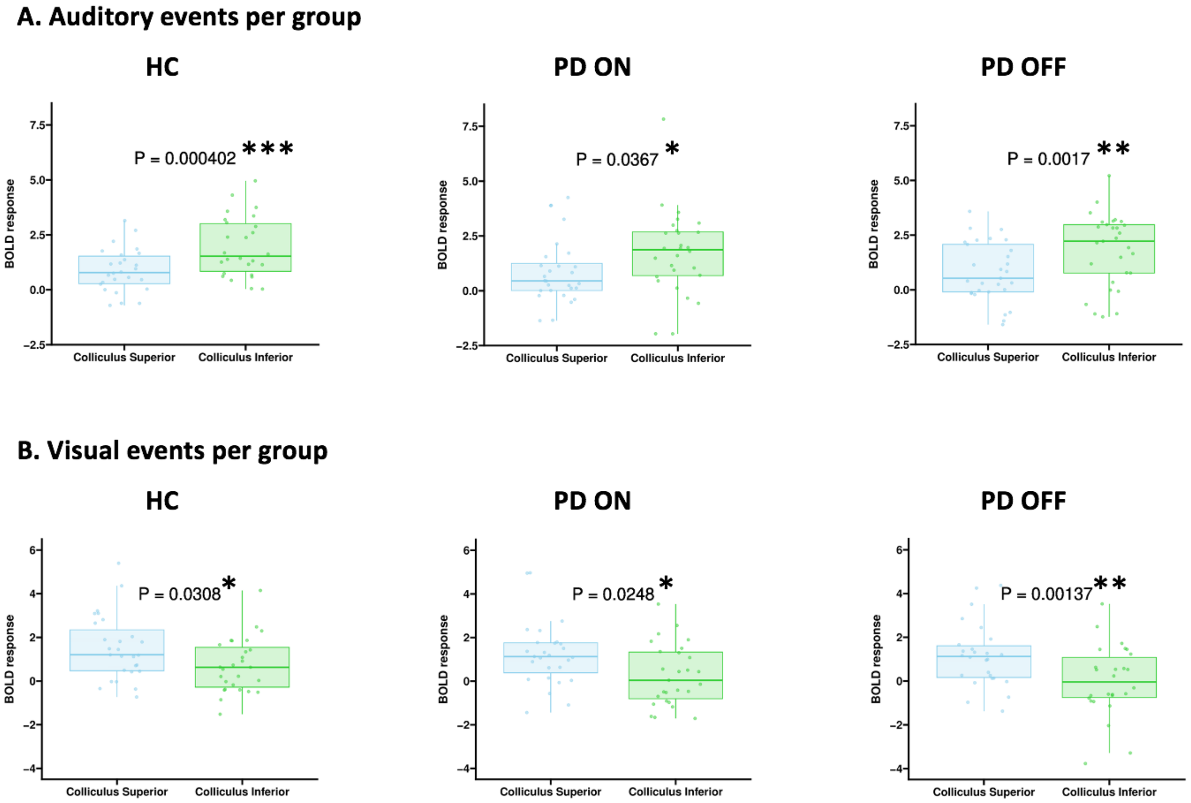


**Supplementary figure 4. BOLD responses for the superior and inferior colliculi during auditory and visual stimuli per group.** Similar to figure 3, but with paired t-tests per group. Shows that all groups show the expected difference. Hence, the lower LC arousal response is unlikely a "gating" problem. Each boxplot shows one data point per subject. *P < 0.05, **P < 0.01, ***P < 0.001.

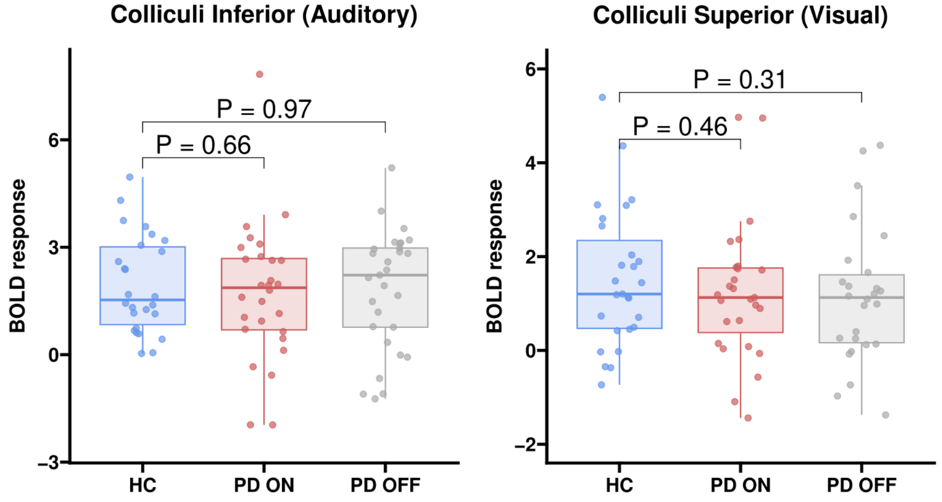


**Supplementary figure 5. Group differences in BOLD responses for the superior and inferior colliculi during auditory and visual stimuli.** Mixed model to test for a group difference in the inferior colliculus to auditory stimuli and the superior colliculus to visual stimuli. There were no group differences (all P > 0.05). Each boxplot shows one data point per subject.

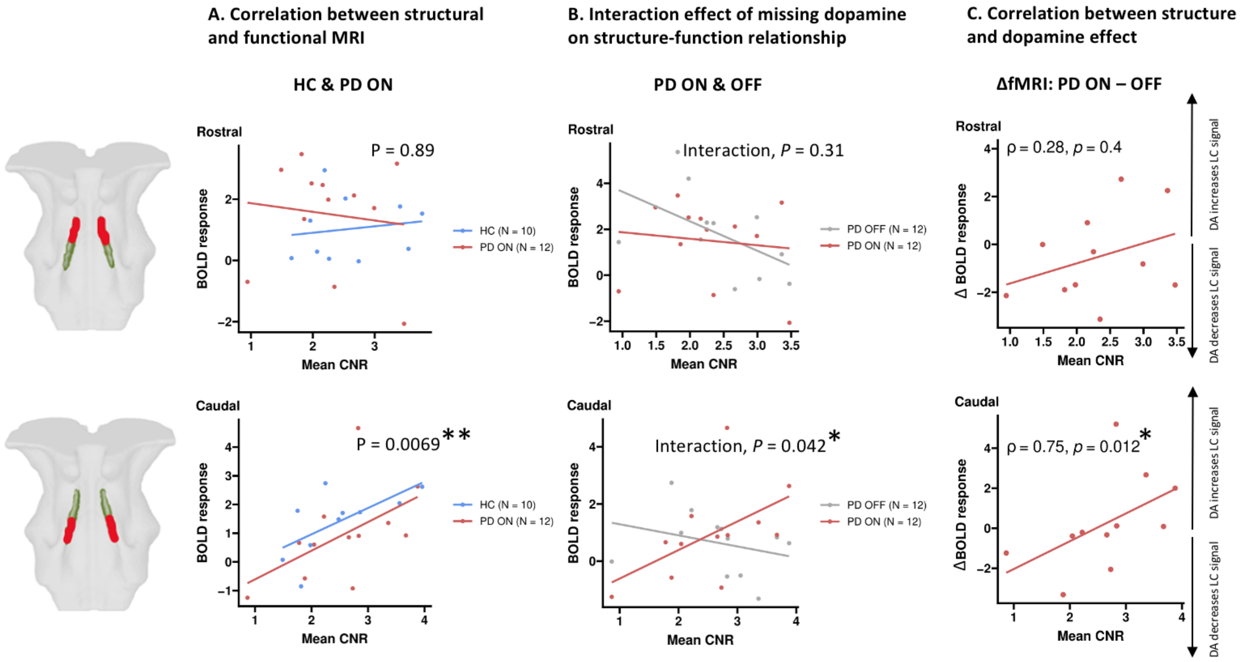


**Supplementary figure 6.** Similar to figure 5, but including the rostral region. *P < 0.05, **P < 0.01, ***P < 0.001.

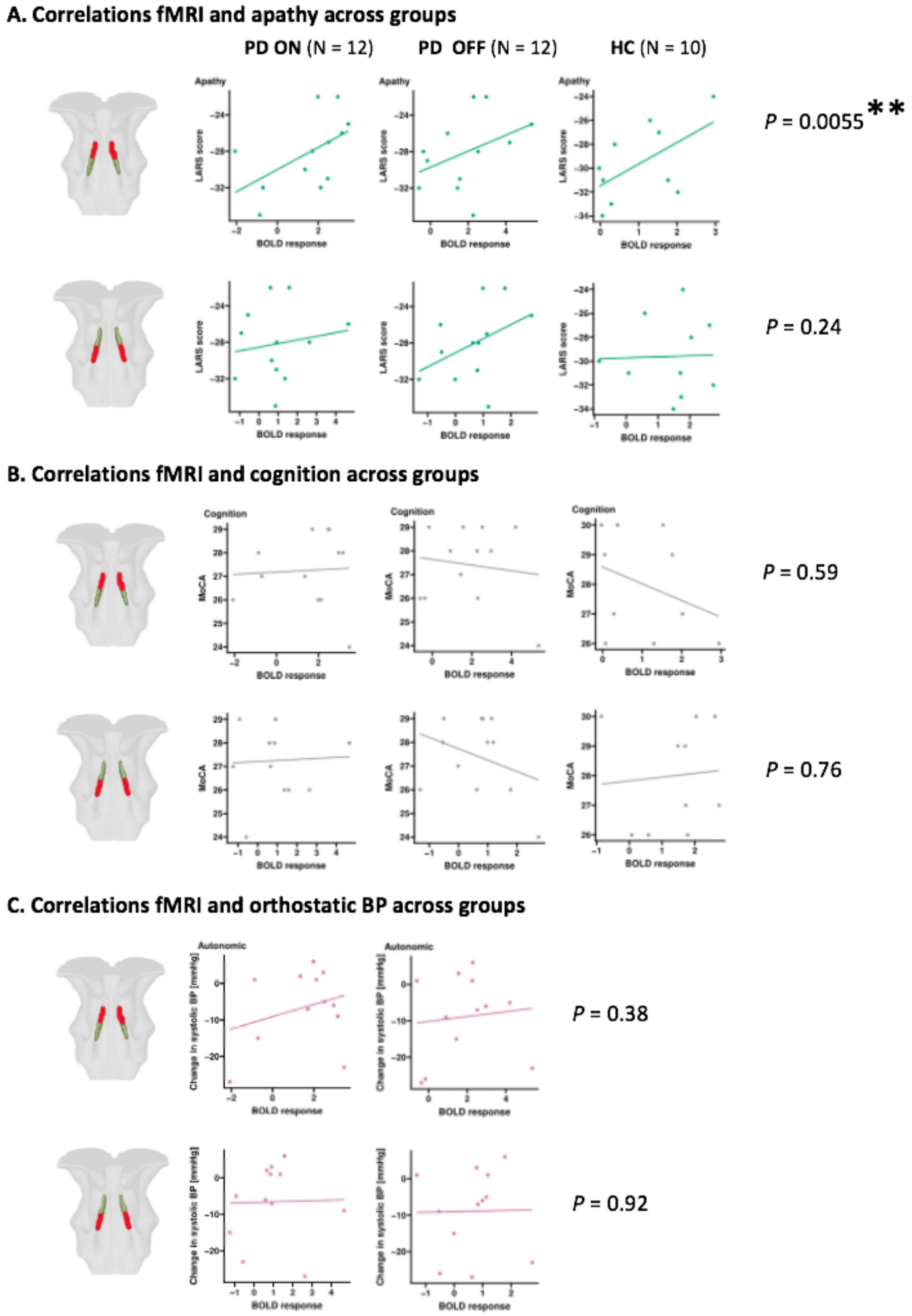


**Supplementary figure 7.** Similar to figure 6, but including MoCA and systolic changes in orthostatic blood pressure. *P < 0.05, **P < 0.01, ***P < 0.001.

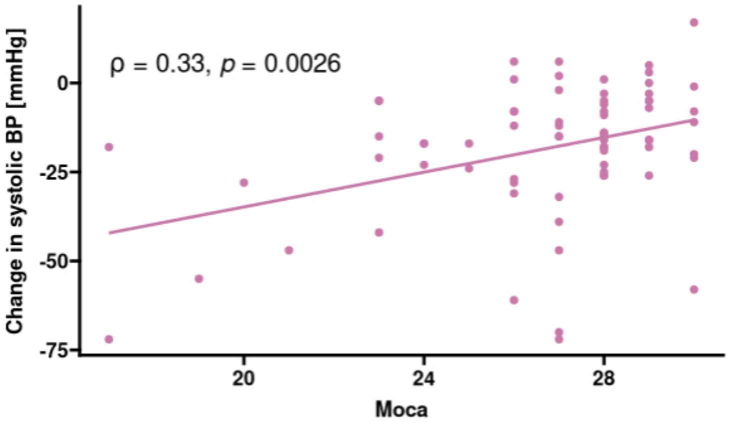


**Supplementary figure 8 correlation between MoCA and orthostatic change in systolic BP.** Spearman correlation showing a significant correlation between MoCA and orthostatic change in systolic BP in patients (rho = 0.33, P = 0.0026). Abbreviations: BP = blood pressure; MoCA = Montreal Cognitive Assessment.

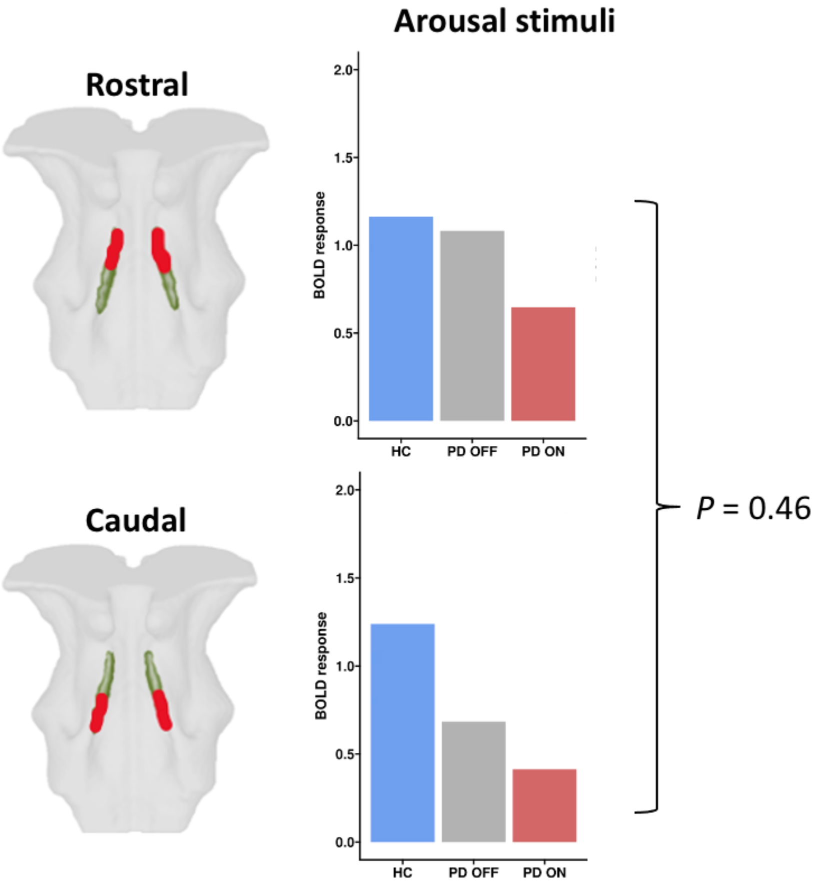


**Supplementary figure 9**. **Rostro-caudal BOLD responses to arousal stimuli.** Difference in mean LC activation between the rostral and caudal regions. There was no difference between rostral and caudal activation (P = 0.46).
